# Sae2 antagonizes Rad9 accumulation at DNA double-strand breaks to attenuate checkpoint signaling and facilitate end resection

**DOI:** 10.1101/424218

**Authors:** Tai-Yuan Yu, Michael Kimble, Lorraine S Symington

## Abstract

The Mre11-Rad50-Xrs2^NBS1^ complex plays important roles in the DNA damage response by activating the Tel1^ATM^ kinase and catalyzing 5’-3’ resection at DNA double-strand breaks (DSBs). To initiate resection, Mre11 endonuclease nicks the 5’ strands at DSB ends in a reaction stimulated by Sae2^CtIP^. Accordingly, Mre11-nuclease deficient (*mre11*-*nd*) and *sae2Δ* mutants are expected to exhibit similar phenotypes; however, we found several notable differences. First, *sae2Δ* cells exhibit greater sensitivity to genotoxins than *mre11*-*nd* cells. Second, *sae2Δ* is synthetic lethal with *sgs1Δ*, whereas the *mre11*-*nd sgs1Δ* mutant is viable. Third, Sae2 attenuates the Tel1-Rad53^CHK2^ checkpoint and antagonizes Rad9^53BP1^ accumulation at DSBs independent of Mre11 nuclease. We show that Sae2 competes with other Tel1 substrates, thus reducing Rad9 binding to chromatin and to Rad53. We suggest that persistent Sae2 binding at DSBs in the *mre11*-*nd* mutant counteracts the inhibitory effects of Rad9 and Rad53 on Exo1 and Dna2-Sgs1 mediated resection, accounting for the different phenotypes conferred by *mre11*-*nd* and *sae2Δ* mutations. Collectively, these data show a resection initiation independent role for Sae2 at DSBs by modulating the DNA damage checkpoint.

## INTRODUCTION

Genomic integrity is constantly threatened by DNA damage that can result from exposure to exogenous sources, such as ionizing radiation, as well as from endogenous sources, including DNA replication errors and intermediates in excision repair or topoisomerase transactions. Cells respond to these insults by an elaborate network of surveillance mechanisms and DNA repair pathways, referred to as the DNA damage response (DDR) (1). DNA double-strand breaks (DSBs) are one of the most cytotoxic forms of DNA damage and can cause loss of genetic information, gross chromosome rearrangements or even cell death in the absence of the appropriate response.

Typically, cells repair DSBs by non-homologous end joining (NHEJ) or by homologous recombination (HR) (2). The Ku heterodimer (Ku70-Ku80), an essential NHEJ component, binds to DSB ends to protect them from degradation, and recruits the DNA ligase IV complex to catalyze end ligation (3). HR employs extensive homology and templated DNA synthesis to restore the broken chromosome, and is considered to be a mostly error-free mode of repair (4). HR initiates by nucleolytic degradation of DNA ends to generate long 3’ single-strand DNA (ssDNA) tails, a process termed end resection. RPA initially binds to the 3’ overhangs to protect them from degradation (5), and is subsequently replaced by Rad51 to promote homologous pairing and strand invasion (6). Initiation of end resection is activated during S and G2 phases of the cell cycle when the sister chromatid is available as a repair template, and is considered to be the main regulatory step in repair pathway choice (7-9). In coordination with DNA repair mechanisms, cells respond to DSBs by a signaling cascade to halt cell cycle progression, induce transcription and to activate key repair proteins (1). Tel1^ATM^ and Mec1^ATR^ are the sentinel kinases that respond to DSBs in *Saccharomyces cerevisiae* (1, 10). Mre11-Rad50-Xrs2^Nbs1^ (MRX^N^) complex bound to ends recruits and activates Tel1^ATM^, whereas Mec1-Ddc2^ATR-ATRIP^ binds to RPA-coated ssDNA generated by end resection. Yeast Tel1 and Mec1 redundantly phosphorylate multiple DNA repair proteins, as well as the downstream effector kinase, Rad53^CHK2^. Rad53 phosphorylation requires the Rad9^53BP1^ adaptor protein, which is recruited to chromatin by Dot1-methylated histone H3-K79, phosphorylated H2A^H2AX^ (*γ*H2A), the 9-1-1 damage clamp and Dpb11^TOPBP1^ (1).

In addition to activating Tel1 kinase, MRX plays critical roles in tethering DNA ends and initiating end resection (4). The current model for end resection is for the Mre11 endonuclease to nick the 5’ strands internal to the DSB ends in a reaction stimulated by cyclin dependent kinase (CDK)-phosphorylated Sae2^CtIP^ (Fig 1A) (11-14). Mre11 exonuclease then degrades in the 3’-5’ direction toward the break ends while more extensive processing of the 5’ strands is catalyzed by Exo1 or by Sgs1 helicase in concert with Dna2 endonuclease (15, 16). MRX also plays a non-catalytic role in end resection by recruiting Dna2 and Sgs1 to DSBs (17). In budding yeast, resection initiation by Mre11 nuclease and Sae2 is essential to remove covalently bound proteins, such as Spo11 from meiotic DSBs, and hairpin-capped DNA ends, but is not essential for processing ends generated by endonucleases (16). In the absence of Mre11 nuclease (e.g., *mre11*-*H125N* mutant) or Sae2, resection of endonuclease-induced DSBs occurs primarily through the activity of Dna2-Sgs1. Thus, the *mre11*-*H125N sgs1Δ* double mutant exhibits greatly increased DNA damage sensitivity and delayed resection of an HO endonuclease-induced DSB relative to the single mutants, while the *sae2Δ sgs1Δ* double mutant is inviable (17-19). Exo1 can only efficiently promote resection at DNA ends when Ku is eliminated from cells. Consequently, deletion of *YKU70* or *YKU80* (encoding the Ku heterodimer) suppresses DNA damage sensitivity and end resection defects of *mre11Δ*, *mre11*-*H125N* and *sae2Δ/ctp1Δ* mutants, and also bypasses lethality of *sae2Δ sgs1Δ* cells, in an Exo1-dependent fashion (17, 18, 20-22).

**Figure 1.**
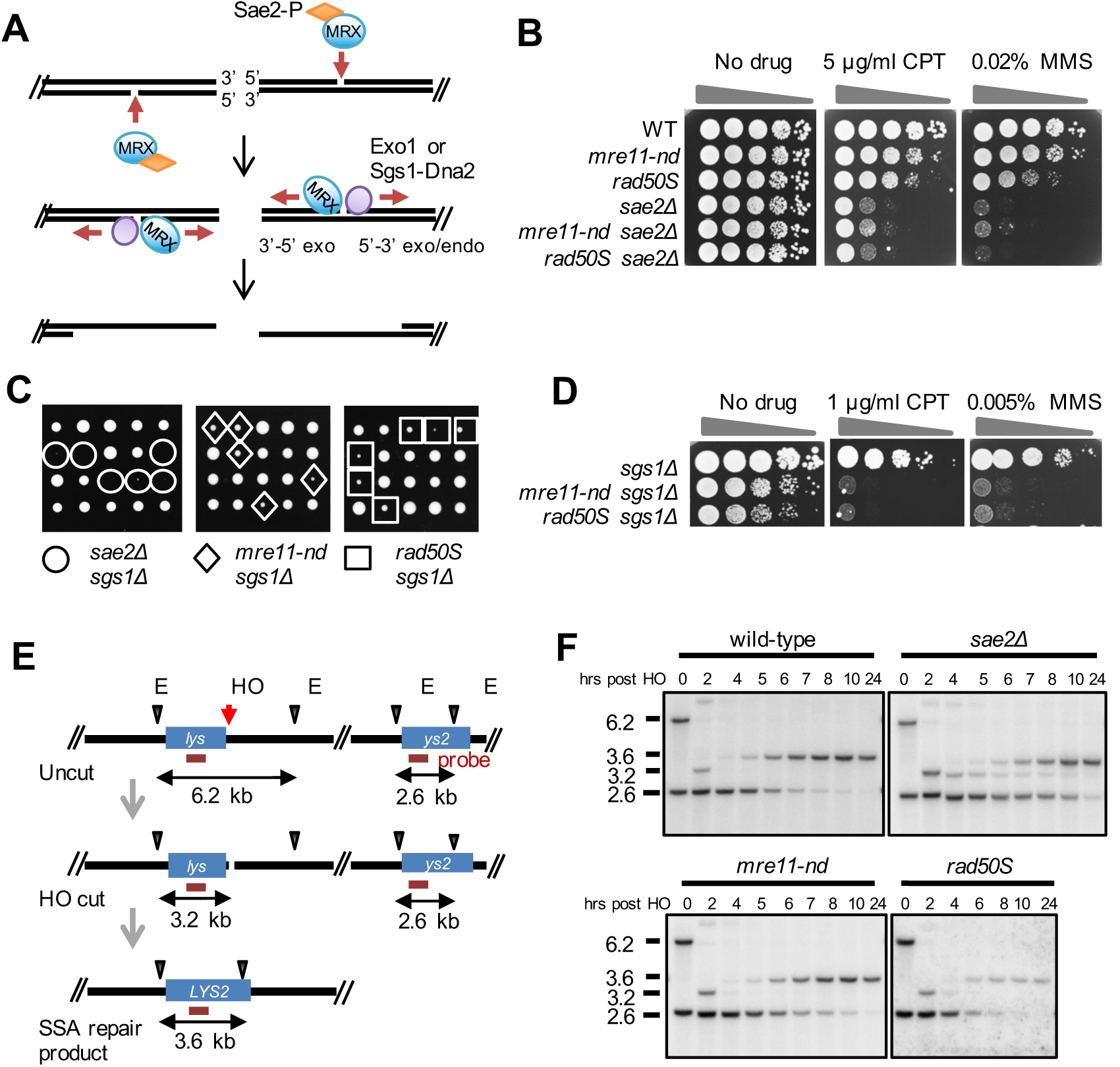
*mre11*-*nd*, *sae2Δ* and *rad50S* mutants display different phenotypes. (A) Model for resection initiation at DSBs (see text for details). (B,D) Ten-fold serial dilutions of the indicated strains spotted on plates without drug, or plates containing camptothecin (CPT), or methyl methanesulfonate (MMS) at the indicated concentrations. (C) *SAE2/sae2Δ* (or *MRE11/mre11*-*nd* or *RAD50/rad50S*) *sgs1Δ/SGS1* heterozygous diploids were sporulated and tetrads were dissected on YPD plates. (E) Schematic of SSA assay showing location of the HO cut site and EcoRV (E) sites used to monitor DSB formation and deletion product. The horizontal red line indicated the sequences to which the probe hybridizes. (F) EcoRV-digested genomic DNA from the indicated strains, before and after HO induction was separated by agarose-gel electrophoresis, blotted and hybridized with a fragment internal to the *LYS2* gene.

In contrast to the *mre11Δ* mutant, which exhibits reduced Mec1 signaling in response to an un-repairable DSB due to resection defects, *mre11*-*H125N* and *sae2Δ* mutations confer normal Mec1 activation (23). Furthermore, phosphorylated Rad53 persists in *sae2Δ* cells, and over-expression of Sae2 diminishes Rad53 activation even though resection is unaffected, suggesting a role for Sae2 in attenuating DNA damage signaling (23). Mre11 persists at DSBs in *sae2Δ* cells due to a defect in end clipping, resulting in hyper-activation of the Tel1 checkpoint and suppression of *mec1Δ* sensitivity to genotoxic agents that cause replication-fork stalling (24-26). Consistent with findings in budding yeast, deletion of *ctp1* suppresses fission yeast *rad3Δ* DNA damage sensitivity by hyper-activation of the Tel1-Chk1 checkpoint (27).

In efforts to further understand the role of Sae2 in the DNA damage response, several groups identified suppressor mutations that bypass *sae2Δ* camptothecin (CPT) sensitivity (28-31). One class of suppressors consists of point mutations within the N-terminal domain of Mre11 that broadly suppress *sae2Δ* sensitivity to a variety of DNA damaging agents (29-31). The *mre11*-*H37Y* and *mre11*-*P110L* mutations, which were characterized in detail, encode proteins with reduced DNA binding affinity and suppress Mre11 hyper-accumulation at DNA ends in the *sae2Δ* background resulting in reduced checkpoint signaling. Another class of suppressors includes components of the DNA damage checkpoint. Rad9 accumulates close to DSBs in *sae2Δ* cells, and elimination of Rad9 restores Sgs1-Dna2 dependent end resection and DNA damage resistance to *sae2Δ* cells (32, 33). Additional *sae2Δ* suppressors include a gain of function *SGS1* allele that overcomes the Rad9 inhibition to end resection, and *tel1* and *rad53* point mutations that reduce Rad9 binding, dampen checkpoint signaling and increase Dna2-Sgs1 dependent resection (28).

The goal of the current study was to investigate the differential sensitivity of *sae2Δ* and *mre11*-*H125N* mutants to DNA damaging agents. We provide evidence that Sae2 counteracts Rad9 binding to DSBs, independent of resection initiation, and the hyper-accumulation of Rad9 in the absence of Sae2 increases Rad53 activation and inhibits resection by Dna2-Sgs1 and Exo1.

## RESULTS

### Differences between Sae2 and Mre11 nuclease-deficient cells

By the current model for resection initiation (Fig 1A), the Mre11 nuclease defective (*mre11-H125N*, hereafter referred to as *mre11*-*nd*) and *sae2Δ* mutants should exhibit similar DNA damage sensitivity. However, we find the *sae2Δ* mutant to be more sensitive to CPT and methyl methanesulfonate (MMS) than the *mre11*-*nd* mutant, and the double mutant exhibits the same sensitivity as the *sae2Δ* single mutant (Fig 1B). Furthermore, *sae2Δ* is lethal when combined with *sgs1Δ*, whereas the *mre11*-*nd sgs1Δ* double mutant is viable (Fig 1C). The *rad50*-*K81I* (*rad50S*) mutant, which is also defective for Spo11 removal from meiotic DSBs and Mre11-catalyzed end clipping in vitro (12, 34), exhibits DNA damage sensitivity intermediate between *mre11*-*nd* and *sae2Δ*. The *rad50S sgs1Δ* double mutant is viable, but shows reduced proliferation and decreased DNA damage resistance relative to the single mutants, similar to *mre11*-*nd sgs1Δ* (Fig 1C, D).

We measured end resection in the *mre11*-*nd*, *rad50S* and *sae2Δ* mutants by a single-strand annealing (SSA) assay. A strain was constructed with two fragments of the *lys2* gene, which share 2.2 kb homology, separated by 20 kb on chromosome V (Fig 1E). An HO endonuclease cut site was inserted at the junction of one *lys2* repeat and the intervening sequence. Following DSB induction, the single-stranded regions of *lys2* exposed by end resection anneal to restore *LYS2*, accompanied by deletion of the intervening sequence. Because there are no essential genes in the region deleted, the SSA product is viable. Additionally, the strains contain a *rad51Δ* mutation to prevent repair by break-induced replication, and a galactose-inducible *HO* gene. SSA was monitored by genomic blot hybridization. The HO cut fragments persist for longer in *sae2Δ* cells than observed in wild type (WT), indicative of delayed resection initiation, and deletion product formation is similarly delayed (Fig 1F). By contrast, *mre11*-*nd* and *rad50S* cells show similar kinetics of repair to WT. Despite the differences in timing of product formation, all of the strains exhibit similar survival on galactose-containing medium (Fig S1A). Some previous studies have shown decreased survival of *sae2Δ* cells by SSA using a different strain background (YMV80) (35); the reason for this difference is currently unknown but could be due to relative usage of Exo1 and Dna2-Sgs1. We find SSA to be highly dependent on Exo1 in W303 (Fig S1B), whereas Dna2-Sgs1 is the primary mechanism for extensive resection in YMV80-derived strains (36). The delayed resection initiation in the *sae2Δ* mutant correlates with the increased sensitivity to CPT and MMS, and synthetic lethality with *sgs1Δ*, as compared to *mre11*-*nd* and *rad50S*.

### Accumulation of Rad9 at DSBs is suppressed by Sae2 and is independent of Mre11 nuclease activity

Previous studies have shown persistent Mre11 and Rad9 binding close to DSBs in the *sae2Δ* mutant leading to hyper-activation of Rad53 (23-25, 32, 33). Mre11 binding to DSBs is also increased in the absence of its nuclease activity (25). We compared Mre11, Tel1 and Rad9 binding in response to a single HO endonuclease-induced DSB by chromatin immunoprecipitation (ChIP) in *mre11*-*nd*, *rad50S and sae2Δ* cells. While *sae2Δ* and *rad50S* cells show increased enrichment of Mre11, Tel1 and Rad9 close to the DSB, Rad9 accumulation is the same or lower than WT in *mre11*-*nd* cells, even though Mre11 and Tel1 levels are increased (Fig 2A). Interestingly, Tel1 is increased to even higher levels in the *rad50S* mutant than observed in *sae2Δ* and *mre11*-*nd* mutants, and correlates with longer telomeres (Fig. S2A)

**Figure 2.**
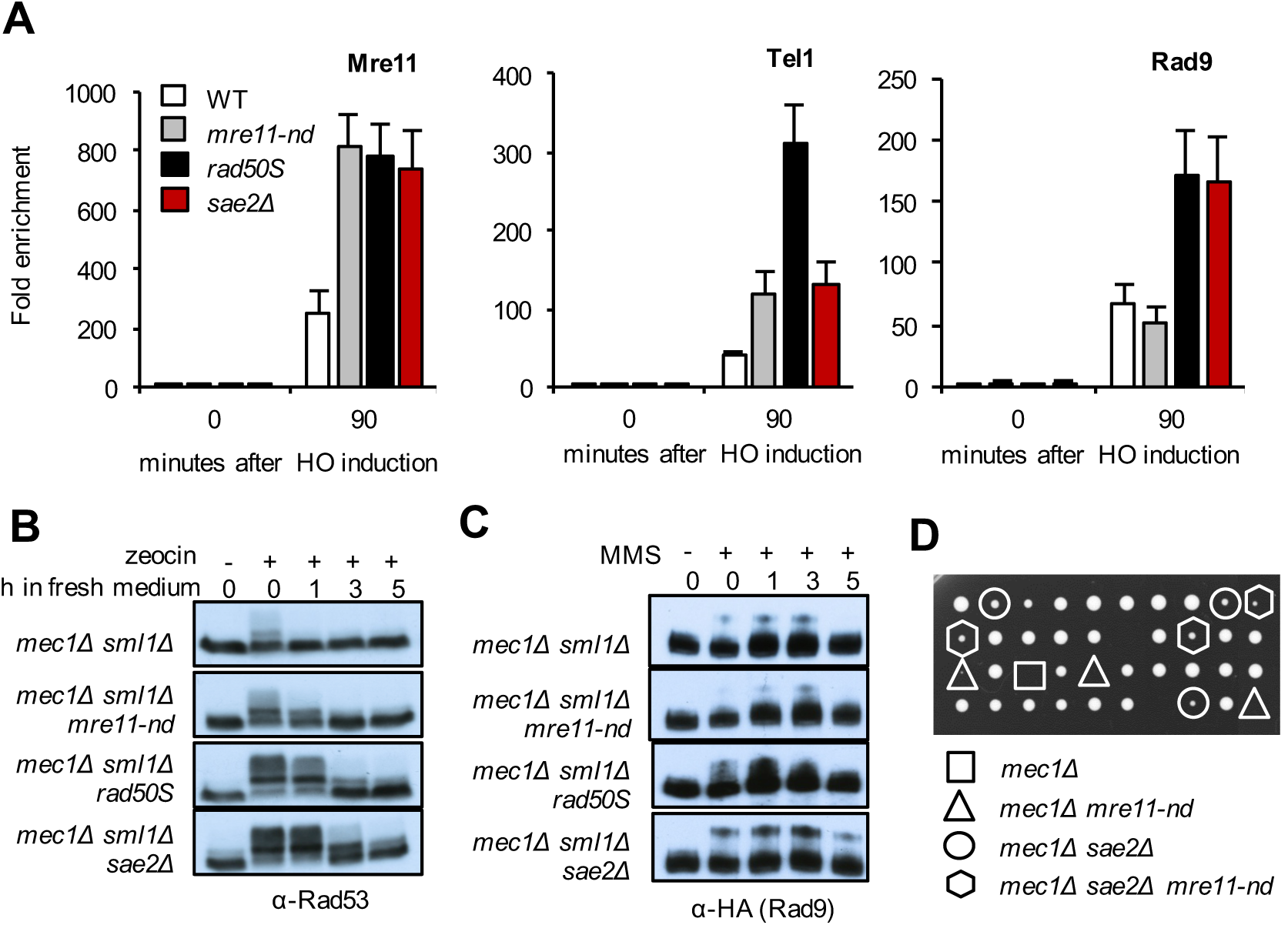
Activation of the Tel1 checkpoint in *sae2Δ* and *rad50S* mutants. (A) The relative fold enrichment of Mre11, HA-Tel1 and Rad9-HA at 0.2 kb from the HO site was evaluated by qPCR after ChIP with anti-Mre11 and anti-HA antibodies. The error bars in all graphs indicate standard deviation (SD) from three biological replicas. (B) Log phase cells (t=0) from the indicated strains were treated with 30 μg/ml Zeocin for 1 hour (h) and released into fresh YPD (t=0-5). Protein samples from different time points before and after drug treatment were analyzed using anti-Rad53 antibodies. (C) Log phase cells (t=0) from the indicated strains were treated with 0.015% MMS for 1 h, released into fresh YPD (t=0-5) and protein samples from different time points before and after drug treatment were analyzed using anti-HA antibodies. (E) A *SAE2/sae2Δ MRE11/mre11*-*nd mec1Δ/MEC1 sml1Δ/SML1* heterozygous diploid was sporulated and tetrads were dissected on YPD plates.

We anticipated that increased Tel1 and Rad9 binding to DSBs in *rad50S* and *sae2Δ* cells would result in enhanced activation of Rad53 phosphorylation relative to WT and *mre11*-*nd* cells. To specifically detect Tel1 activation, we monitored Rad53 phosphorylation in response to transient zeocin treatment of Mec1-deficient cells (all strains included *sml1Δ* to suppress lethality of *mec1Δ*). Rad53 activation and recovery are similar in *mec1Δ* and *mec1Δ mre11*-*nd* cells, whereas *mec1Δ rad50S* and *mec1Δ sae2Δ* cells show increased Rad53 phosphorylation and delayed recovery (Fig 2B), in agreement with a previous study (24). Similar responses were observed following treatment of cells with MMS or CPT (Fig S2B). Consistent with Rad53 activation, Rad9 phosphorylation is increased in *mec1Δ rad50S* and *mec1Δ sae2Δ* mutants in response to MMS, as compared to WT and *mre11*-*nd* cells (Fig 2C). Moreover, *sae2Δ* suppresses lethality of *mec1Δ SML1* cells in a Tel1 and Rad9 dependent manner, while *mre11*-*nd* and *rad50S* fail to suppress *mec1Δ* lethality (Fig 2D, S2C, S2D). Even though *rad50S* cells exhibit greater enrichment of Tel1 at DSBs than observed in *sae2Δ* cells, Rad53 activation is higher in the absence of Sae2 and this could account for suppression of *mec1Δ* lethality.

### Rad9 accumulation at DSBs and Rad53-catalyzed phosphorylation of Exo1 contribute to *sae2Δ* DNA damage sensitivity

In agreement with published studies, we found that deletion of *RAD9* suppresses the CPT and MMS sensitivity of the *sae2Δ* mutant (Fig 3A) (28, 32). Surprisingly, *rad9Δ* had no suppressive effect in the *mre11*-*nd* background, and even resulted in higher sensitivity to CPT and MMS than the single mutants (Fig 3A). Because the increased DNA damage sensitivity of *sae2Δ* cells, as compared to *mre11*-*nd*, appears to result from Rad9 accumulation, we anticipated that by decreasing Rad9 binding, we should restore damage resistance to *sae2Δ* cells. Recruitment of Rad9 to chromatin requires phosphorylation of H2A-S129 by Te11 and/or Mec1, and Dot1-methylated H3-K79 (37-39). Consistently, we observe higher H2A phosphorylation (*γ*H2A) close to the HO-induced DSB in *sae2Δ* cells compared with *mre11*-*nd* or WT (Fig S3A). We anticipated that *dot1Δ* and *hta1*-*S129A* mutations would partially suppress *sae2Δ* DNA damage sensitivity because both have been shown to increase DNA end resection and the *hta*-*S129A* mutation was shown to partially suppress the end resection defect of *sae2Δ* cells (32, 33, 40-42). Surprisingly, the *hta*-*S129A* mutation fails to suppress *sae2Δ* DNA damage sensitivity (Fig S3B) (32, 33). The slight synergism between *sae2Δ* and *hta*-*S129A* mutations could be due to the role of *γ*H2A in stabilizing replications forks and/or sister-chromatid recombination coupled to the resection defect (43, 44). By contrast, *dot1Δ* partially suppresses *sae2Δ* DNA damage sensitivity. In addition to the nucleosome-dependent pathway for recruitment, Rad9 binds to Dpb11^TopBP1^, which is tethered to damage sites by the 9-1-1 clamp (Fig 3B) (45). The Rad9 S462A and T474A mutations (hereafter referred to as *rad9*-*2A*) eliminate the interaction between Rad9 and Dpb11 (46). A previous study showed that the *rad9*-*2A* mutation suppresses the *sae2Δ* SSA defect (32); consistently, we find the *rad9*-*2A* mutation restores DNA damage resistance to *sae2Δ* cells (Fig 3A).

**Figure 3.**
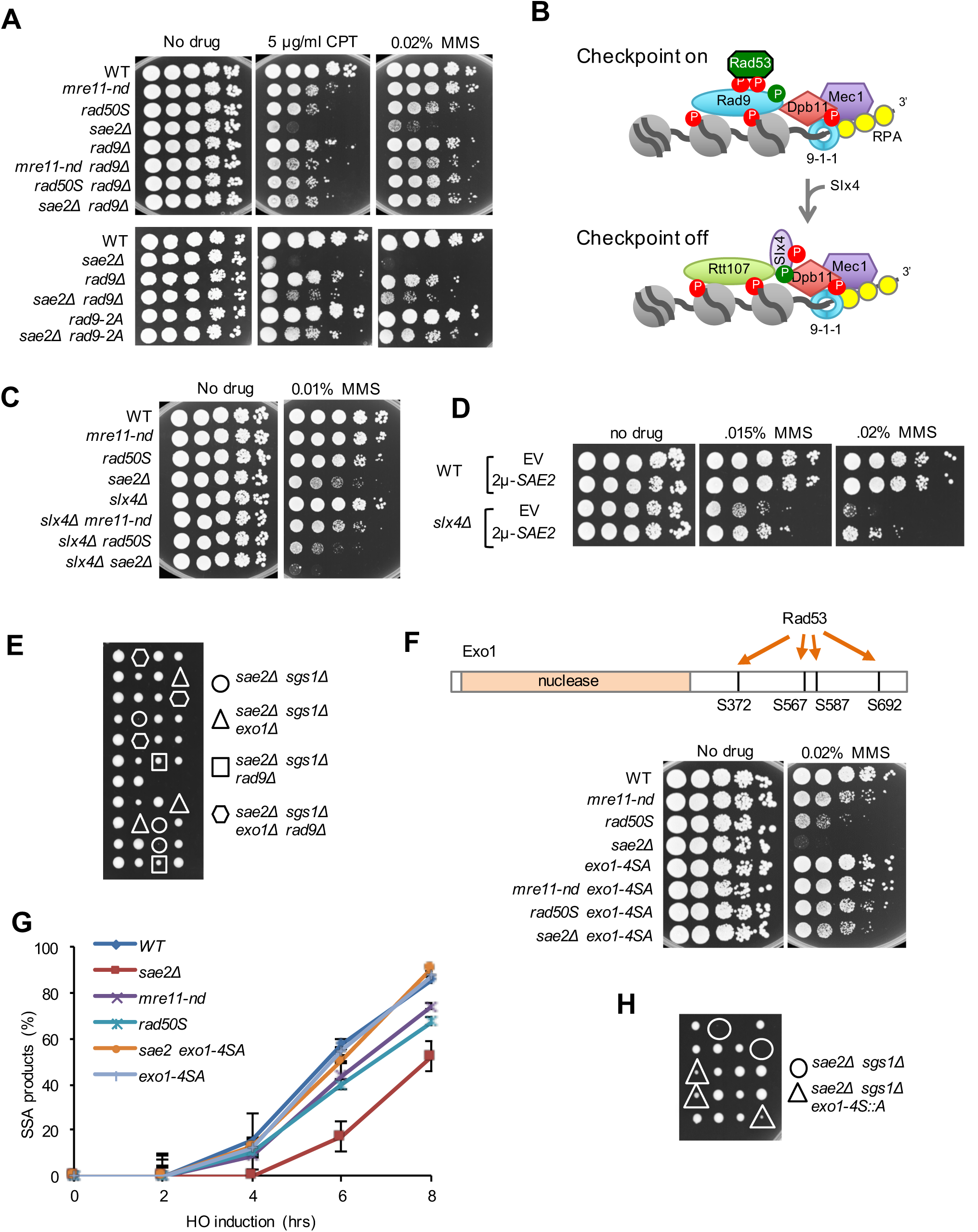
Rad9 chromatin binding and Rad53 activation contribute to *sae2Δ* DNA damage sensitivity and *sae2Δ sgs1Δ* lethality. (A) Ten-fold serial dilutions of the indicated strains spotted on plates without drug, or plates containing indicated DNA damaging agents. (B) Schematic showing stabilization of Rad9 binding to chromatin by Dpb11 to activate the DNA damage checkpoint. CDK (green circles) and Mec1 (red circles) phosphorylated Slx4 competes with Rad9 for Dpb11 interaction, dampening the checkpoint. (C) Ten-fold serial dilutions of the indicated strains spotted on plates without drug, or plates containing indicated DNA damaging agents. (D) Ten-fold serial dilutions of the indicated strains spotted on plates without drug, or plates containing MMS. (E) Diploids heterozygous for the indicated mutations were sporulated and tetrads dissected on YPD plates. (F) Upper panel schematic showing Rad53-dependent phosphorylation sites in the C-terminal region of Exo1, lower panel shows Ten-fold serial dilutions of the indicated strains spotted on plates without drug, or plates containing indicated DNA damaging agents. (G) SSA kinetics were assessed by qPCR of genomic DNA from the indicated strains before and after HO induction. Error bars indicated SD from three independent trials. (H) Diploids heterozygous for the indicated mutations were sporulated and tetrads dissected on YPD plates.

Slx4 and its binding partner, Rtt107, compete with Rad9 for interaction with Dpb11 and *γ*H2A, dampening DNA damage signaling (47). Since our studies indicate that Sae2 antagonizes Rad9 accumulation at DSBs, we tested genetic interaction between *sae2Δ* and *slx4Δ*. Consistent with a previous study *slx4Δ* synergizes with *sae2Δ* (48), and also with *rad50S* and *mre11*-*nd* (Fig 3C). Moreover, expression of *SAE2* from a high-copy-number plasmid suppresses *slx4Δ* MMS sensitivity, suggesting that Sae2 can substitute for the checkpoint attenuation function of Slx4 (Fig 3D).

Like *rad9Δ*, the *tel1*-*kd* mutation suppresses *sae2Δ* DNA damage sensitivity (33), but fails to suppress the CPT and MMS sensitivity of the *mre11*-*nd* mutant (Fig S3C). Elimination of Rad53 kinase activity, or Rad53 interaction with Rad9 (*rad53-R605A)*, also results in suppression of *sae2Δ* CPT sensitivity (Fig S3D). Notably, elimination of Rad9, or Tel1 or Rad53 kinase activity, equalizes the DNA damage sensitivity of the *mre11*-*nd*, *rad50S* and *sae2Δ* mutants indicating that hyper-activation of the DNA damage checkpoint is responsible for the difference between *sae2Δ* and *mre11*-*nd* mutants.

Previous studies showed that *rad9Δ* suppression of the DNA damage sensitivity and resection defects of *sae2Δ* is by activation of Dna2-Sgs1, and not Exo1 (33). However, the finding that *rad9Δ* suppresses *sae2Δ sgs1Δ* lethality indicates that Exo1 must be activated. Indeed, we find that *rad9Δ* suppression of *sae2Δ sgs1Δ* lethality requires *EXO1* (Fig 3E). Exo1 has been identified as a substrate for the Rad53 kinase (49), and substitution of four serine residues in the C-terminal domain with alanine was shown to increase Exo1 activity at telomeres (50). To determine whether Rad53-catalyzed phosphorylation of Exo1 contributes to the down regulation of resection observed in *sae2Δ* cells, we investigated genetic interaction between *sae2Δ* and *exo1*-*4S::A*. The *exo1*-*4S::A* allele suppresses *sae2Δ* CPT and MMS sensitivity, and the *sae2Δ sgs1Δ* synthetic lethality (Fig 3F, H). The suppressive effect of *exo1*-*4S::A* is also seen in *mre11*-*nd* and *rad50S* backgrounds. The *exo1*-*4S::A* derivatives show different sensitivities to CPT and MMS, which we attribute to suppression of Dna2-Sgs1-catalyzed resection by Rad9, particularly in *sae2Δ* cells. We measured SSA by Southern blot hybridization and quantitative PCR and found the kinetics to be similar in WT and *exo1*-*4S::A* cells (Fig 3G). Notably, the *sae2Δ exo1*-*4S::A* double mutant exhibits faster resection than *sae2Δ.* Thus, the DNA damage checkpoint inhibits resection in Sae2-deficient cells by Rad9 blockade of Dna2-Sgs1 and inhibitory phosphorylation of Exo1 by Rad53.

### Over-expression of *SAE2* reduces Rad9 binding at DSBs

A previous study showed reduced Mre11 association with DSBs and attenuation of Rad53 activation when Sae2 is over-expressed (23). Sae2 over-expression (OE) does not diminish end resection suggesting that the inhibitory effect of Sae2 on Rad53 activation is at a step subsequent to Mec1-Ddc2 recruitment to RPA-coated ssDNA (23). To determine whether Sae2 OE reduces Rad9 accumulation at DSBs we inserted the *GAL* promoter upstream of the *SAE2* locus in a strain expressing a Sae2-Myc fusion. Following galactose induction to simultaneously induce HO cleavage and Sae2 expression, Mre11, Tel1 and Rad9 binding to sequences adjacent to the DSB were measured by ChIP. Consistent with the study by Clerici et al (23), the amount of Mre11 bound to DSBs is decreased by ~70% when Sae2 is OE (Fig 4A); Tel1 and Rad9 binding are also significantly reduced. To determine whether reduced accumulation of Rad9 is a consequence of faster turnover of Mre11 at DSBs, we monitored Mre11 and Rad9 association with DSBs when Sae2 is OE in *mre11*-*nd* and *rad50S* backgrounds. Sae2 OE fails to remove Mre11 from ends when resection initiation is compromised; however, Rad9 levels are reduced (Fig 4B). These data indicate separable functions of Sae2 in turnover of MRX at ends and inhibition of Rad9 binding to chromatin. The reduction in Rad9 binding when Sae2 is OE correlates with decreased Rad53 activation in response to the HO-induced DSB (Fig 4C). Sae2 OE results in greatly increased DNA damage sensitivity of *mre11*-*nd* and *rad50S* mutants, indicating a greater reliance on the checkpoint when repair is delayed (Fig S4A). We immunoprecipitated Rad53 from cells to determine whether the reduction in Rad9 binding caused by Sae2 OE prevents Rad53 interaction. As expected, Rad9 was recovered with Rad53 only after DSB induction. However, the Rad53-Rad9 interaction was greatly reduced in cells with Sae2 OE (Fig 4C). Consistent with the negative effect of Sae2 on Rad53-Rad9 interaction, we find increased Rad53-Rad9 binding in *mec1Δ sae2Δ* cells, compared with *mec1Δ* or *mec1Δ mre11*-*nd* cells (Fig 4D).

**Figure 4.**
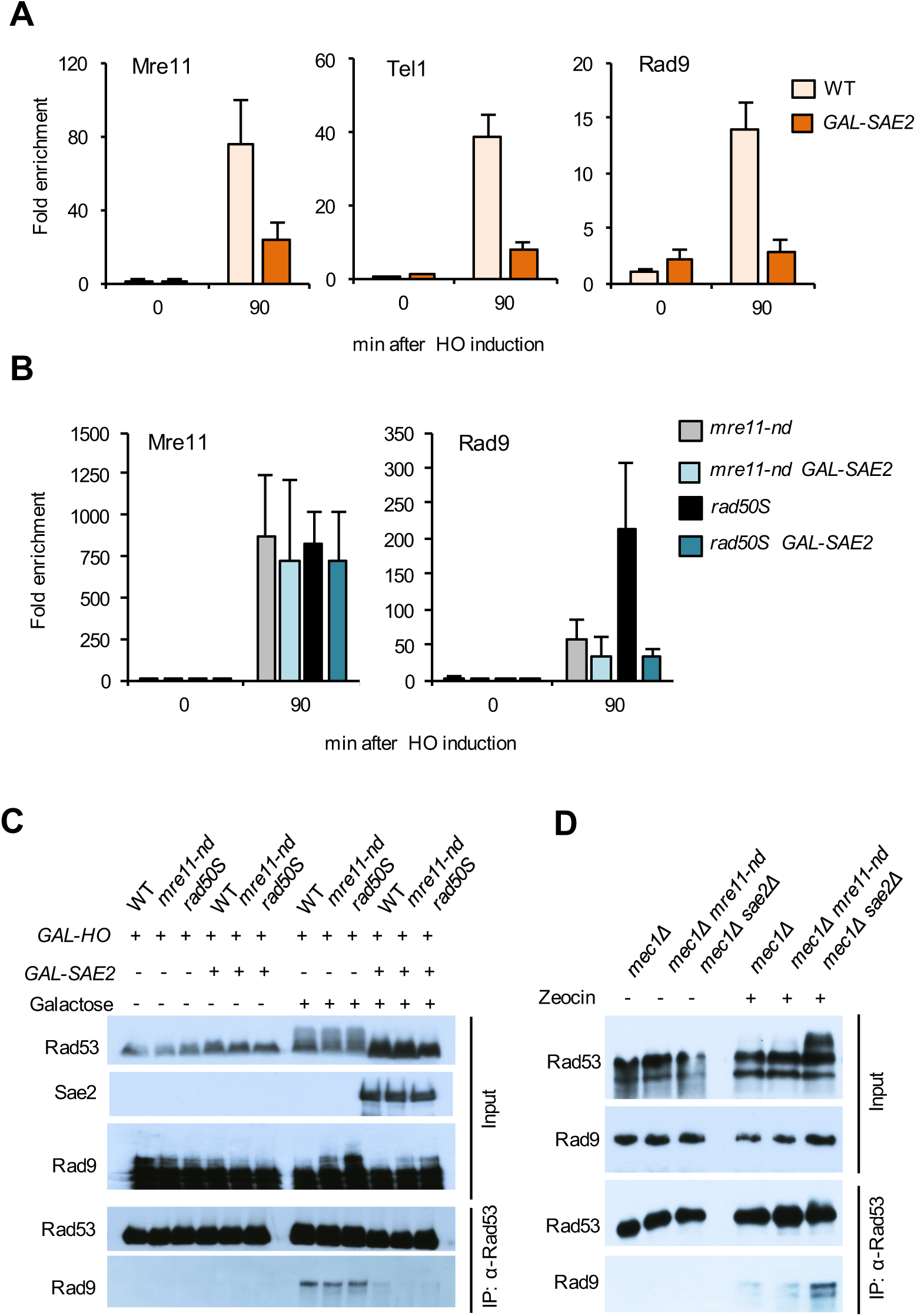
Sae2 OE reduces Rad9 binding at DSBs and to Rad53. (A) Relative fold enrichment of Mre11, Rad9 and Tel1 0.2 kb from the HO cleavage site was evaluated by qPCR after ChIP with anti-Mre11 or anti-HA antibodies. (B) Relative fold enrichment of Mre11 and Rad9 0.2 kb from the HO cut site was measured by qPCR in *mre11*-*nd* and *rad50S* cells with Sae2 OE. (C) Upper panel shows IP inputs and lower panel shows Rad53 immunoprecipitates probed with α-Rad53, Myc (Sae2) or HA (Rad9) antibodies of the indicated strains, before or after galactose induction. (D) Upper panel shows IP inputs and lower panel shows Rad53 immunoprecipitates probed with α-Rad53 or HA (Rad9) antibodies of the indicated strains, before or after MMS treatment. All of the strains in panel D have the *smllΔ* mutation.

### Sae2 phosphorylation by Tel1 and/or Mec1 dampens checkpoint signaling

Mec1 and Tel1 phosphorylate Sae2 on multiple residues in response to DNA damage (Fig 5A), but the physiological role of these modifications has not been firmly established (51, 52). Mutating five of the Mec1/Tel1 sites (S73, T90, S249, T279 and S289) to alanine, *sae2*-*5A*, prevents damage-induced phosphorylation and confers MMS sensitivity (51). A previous study showed that synthetic Sae2 peptides with phosphorylated T90 or T279 residues are able to interact with forkhead-associated domain 1 (FHA1) of Rad53 *in vitro* (53). Moreover, the *sae2*-*T90A*, *T279A* (*sae2*-*2A*) mutant is sensitive to MMS and Rad53 remains hyper-phosphorylated following MMS treatment (53).

**Figure 5.**
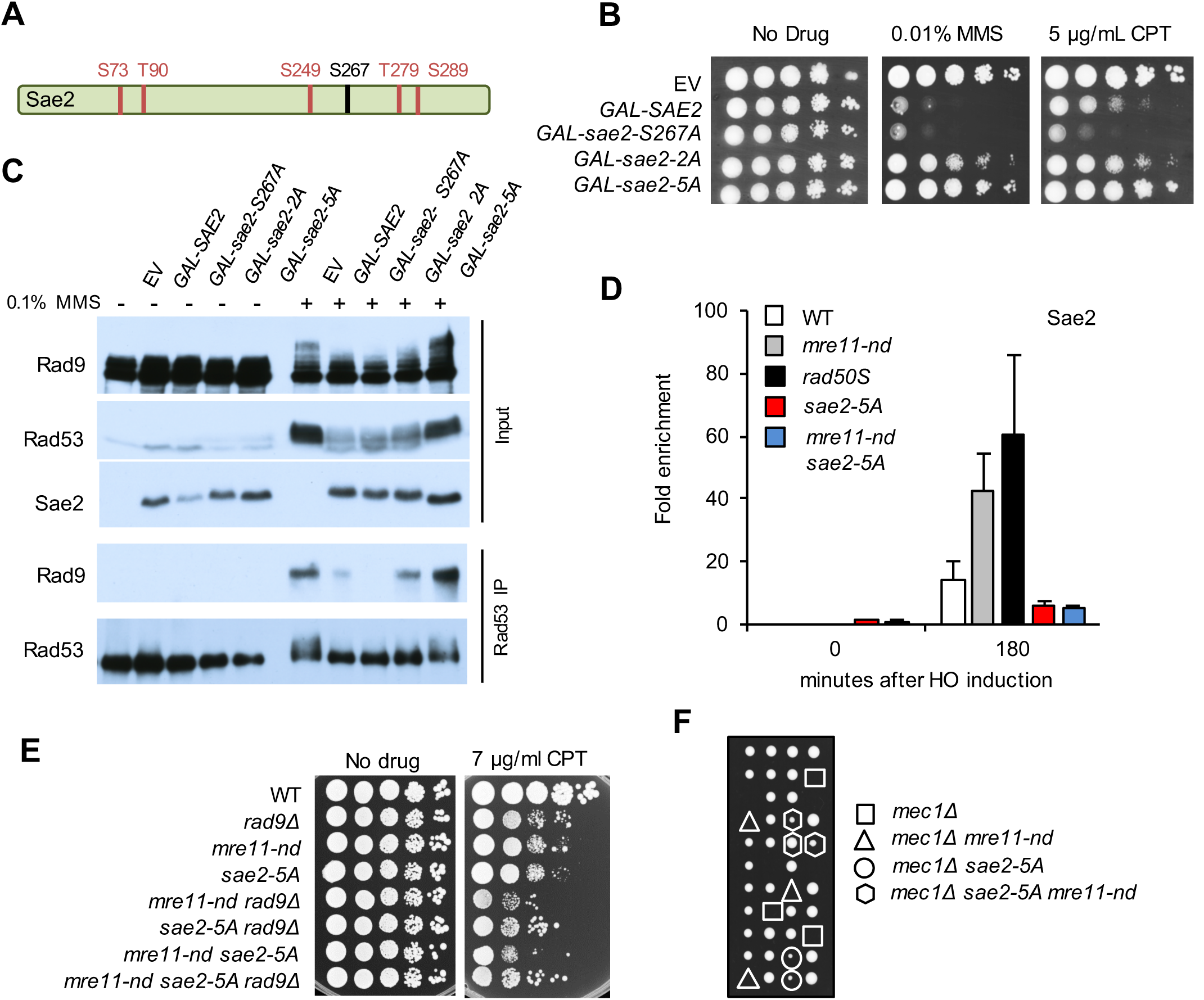
Sae2 attenuates checkpoint signaling by competition for Tel1/Mec1 targets. (A) Schematic of Sae2 showing the main CDK (black) and Mec1/Tel1 (red) phosphorylation sites. (B) Ten-fold serial dilutions of WT cells with *SAE2* or phosphorylation defective *sae2* alleles over-expressed from a *GAL* promoter were spotted on YPGal or YPGal with 0.01% MMS or 5 μg/ml CPT. (C) Western blot of inputs and Rad53 IP following OE of Sae2 or sae2 phosphorylation-site mutants. (D) Relative fold enrichment of Sae2-MYC 0.2 kb from the HO cleavage site was evaluated by qPCR after ChIP with anti-MYC antibodies. (E) Ten-fold serial dilutions of the indicated strains spotted on plates without drug, or plates containing indicated DNA damaging agents. (F) A *SAE2/sae2*-*5A MRE11/mre11*-*nd mec1Δ/MEC1 sml1/SML1* heterozygous diploid was sporulated and tetrads were dissected on YPD plates.

We expressed *SAE2*, *sae2*-*5A* and *sae2*-*2A* alleles from the *GAL* promoter of a centromere-containing plasmid in WT cells. Sae2 OE results in sensitivity to CPT and MMS (Fig 5B), whereas OE of the sae2-2A and sae2-5A variants does not sensitize cells to DNA damage. By contrast, OE of sae2-S267A, which prevents CDK-mediated phosphorylation of Sae2 (14), causes even greater CPT sensitivity than Sae2 OE. The effects of Sae2 OE are similar in WT and *sae2Δ* cells (Fig S4B). Furthermore, OE of Sae2 or sae2-S267A reduces Rad53 and Rad9 phosphorylation in response to MMS, while sae2-5A OE does not (Fig 5C). The reduction in Rad9 phosphorylation correlates with diminished interaction with Rad53, as measured by co-immunoprecipitation (Fig 5C). Thus, Mec1-Tel1 phosphorylation of Sae2 results in attenuation of Rad53 signaling and this is independent of CDK-catalyzed phosphorylation and activation of Mre11 endonuclease.

Because the effects of Sae2 OE are consistent with sequestering Tel1 and/or Mec1 activity, we considered a model whereby a high local concentration of Sae2 when end clipping is abolished by the *mre11*-*nd* or *rad50S* mutation competes with other Te11 substrates, dampening checkpoint activation. In agreement, we find greater enrichment of Sae2 at DSBs in *mre11*-*nd* and *rad50S* cells than observed for WT (Fig 5D). The sae2-5A mutant protein shows decreased chromatin binding, consistent with the impaired checkpoint dampening function of sae2-5A. Furthermore, *sae2*-*5A* synergizes with *mre11*-*nd* for DNA damage sensitivity in a Rad9-dependent manner and suppresses *mec1Δ* lethality (5E, F). These data support the hypothesis that PIKK phosphorylation of Sae2 is required for its role in checkpoint attenuation.

## DISCUSSION

Previous studies demonstrated increased DNA damage sensitivity and a greater delay in resection initiation caused by *sae2Δ* as compared to mutations that inactivate the Mre11 nuclease activity (17, 18, 29, 54), indicating that Sae2 has a function in the DNA damage response that is independent of Mre11 endonuclease activation. We show here that the increased DNA damage sensitivity of *sae2Δ* cells is due to excessive Rad9 binding in the vicinity of DSBs and hyper-activation of the Rad53 checkpoint. While both *sae2Δ* and *mre11*-*nd* cells show elevated levels of Mre11 and Tel1 at DSBs as a consequence of delayed resection initiation, increased Rad9 binding is not seen in *mre11*-*nd* cells. We suggest that the delay in resection initiation caused by *mre11-nd* leads to a high local concentration of Sae2 in the vicinity of a DSB that competes with Rad9 for Tel1 activity, lowering the amount of Rad9 retained at DSBs and reducing Rad53 activation (Fig 6). The increased accumulation of Rad9 at damaged sites in the *sae2Δ* mutant acts as a barrier to Dna2-Sgs1 resection, and hyper-activation of the Rad53 kinase results in inhibitory phosphorylation of Exo1. These two mechanisms contribute to the greater delay in end resection and higher DNA damage sensitivity of *sae2Δ* cells relative to *mre11*-*nd* cells.

**Figure 6.**
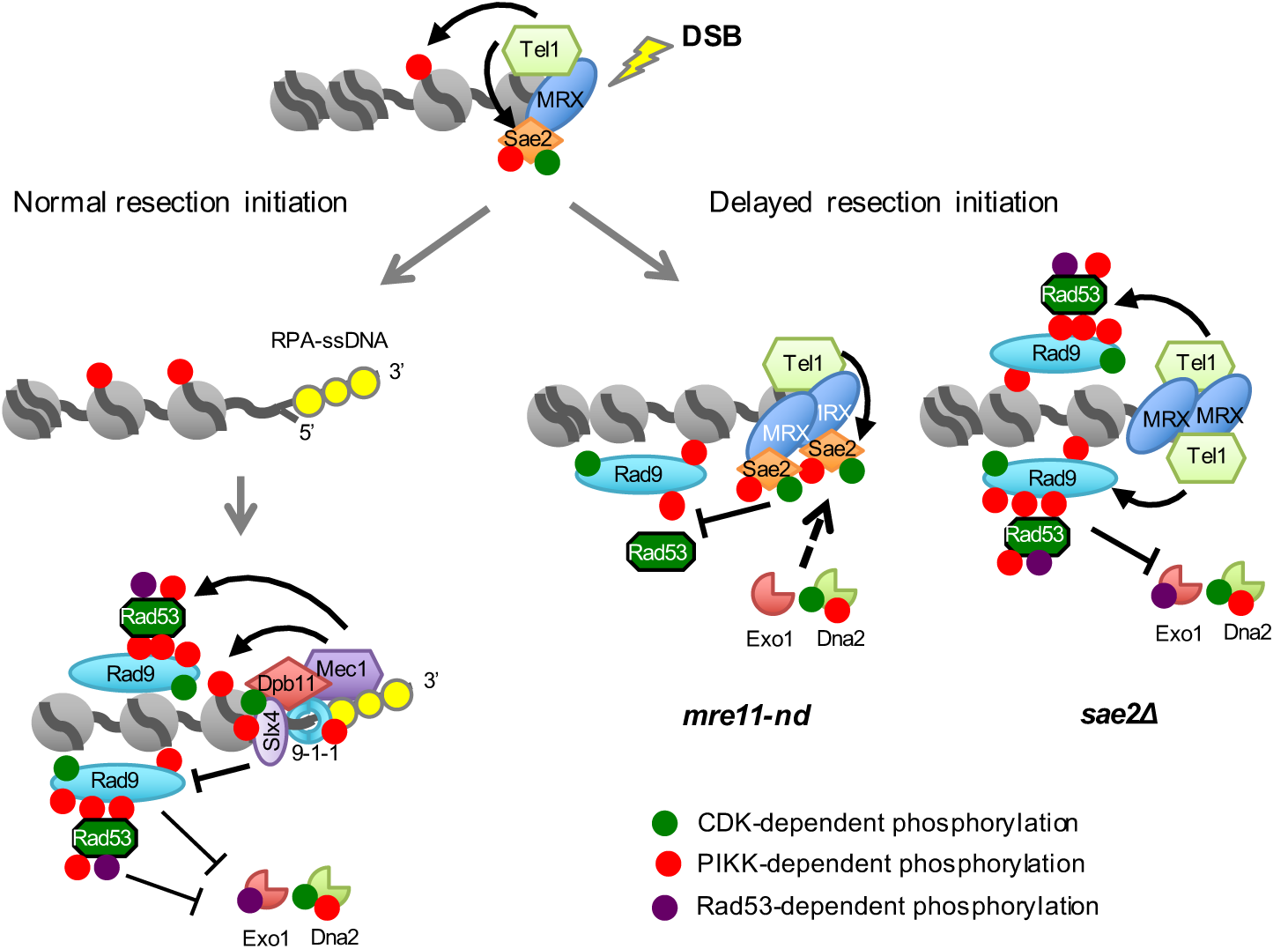
Sae2 controls short and long-range resection pathways. Normally, resection initiation by Mre11 and Sae2 is fast resulting in low dwell time of Tel1 at DSBs, and consequently low activation of Tel1 kinase. After resection initiation, Mec1-Ddc2 is recruited to ssDNA overhangs, phosphorylating Rad9 and Rad53 to slow down resection by Dna2-Sgs1 and Exo1. Slx4-Rtt107 competes with Rad9 for Dpb11 binding to dampen the checkpoint and, consequently, increase extensive resection. When resection initiation is compromised by the *mre11*-*nd* mutation, MRX, Tel1 and Sae2 accumulate at DSBs and Sae2 competes with other Tel1 substrates for phosphorylation, reducing Rad9 binding, Rad53 activation and allowing resection by Dna2-Sgs1 and Exo1. In the absence of Sae2, Tel1 is hyper-activated, causing increased Rad9 binding and Rad53 activation, thereby diminishing resection by Dna2-Sgs1 and Exo1.

The *rad50S* mutant behaves differently to *mre11*-*nd* and *sae2Δ*. While the DNA damage sensitivity and resection defects of *rad50s* cells are similar to *mre11*-*nd*, accumulation of Rad9 at DSBs resembles *sae2Δ*. If Rad9 accumulation is responsible for reduced resection in *sae2Δ* cells, then how do we explain the *rad50S* phenotype? Tel1 binding is much higher in *rad50S* than in *sae2Δ* cells and telomeres are longer (Fig S2A) (55), consistent with Tel1 hyperactivation; however, Rad53 activation (in the *mec1Δ* background) is slightly lower and *rad50S* fails to suppress *mec1Δ* lethality. We suggest that the high local concentration of Sae2 in *rad50S* cells attenuates Rad53 activation and this allows more efficient resection by Exo1. Indeed, the *exo1*-*4SA* mutation is more effective in suppressing *rad50S* DNA damage sensitivity than *rad9Δ*. We suggest that most of the checkpoint dampening role of Sae2 is through inhibition of Rad9 accumulation near DSBs, but cannot rule out additional roles via direct interaction with Rad53 and Dun1 (53).

Rad9 and its mammalian ortholog, 53BP1, have well documented roles in preventing end resection (42, 56). In yeast, the Dna2-Sgs1 resection mechanism is the primary target of Rad9 inhibition (57), particularly in *sae2Δ* cells (28, 32). In the absence of Sae2 and Mre11 endonuclease-catalyzed incision there is no nick for Exo1 entry and Ku is highly effective in preventing Exo1 resection at ends; thus, resection in *mre11*-*nd* cells is mostly dependent on Dna2-Sgs1 (17-19). The increased Rad9 binding in *sae2Δ* as compared to *mre11*-*nd* cells presumably creates a greater barrier for Dna2-Sgs1 accounting the more severe phenotype of the *sae2Δ* mutant. By contrast to budding yeast, the increased resection observed in the absence of 53BP1 in mouse cells is largely dependent on CtIP (58). There are two possible explanations for this difference. First, CtIP might be required to recruit DNA2 to DSBs or function with DNA2-BLM/WRN in long-range resection; this idea is supported by studies showing epistasis between CtIP and DNA2 for resection defects and direct stimulation of DNA2-BLM activity by CtIP (59-62). Second, if 53BP1 recruitment to DSBs occurs prior to resection initiation it could potentially create a barrier to CtIP and MRN-catalyzed incision.

Resection initiation generates RPA-bound ssDNA, which is essential for Mec1-Ddc2 signaling, as well as creating a recessed 5’ end for loading the 9-1-1 damage clamp (1). Previous studies have shown that 9-1-1 has both positive and negative roles in regulating end resection (57), both of which are modulated by the multi-BRCT domain protein, Dpb11^TOPBP1^. Dpb11 interacts with the Ddc1 subunit of the 9-1-1 complex and Rad9 at damage sites to mediate Rad53 phosphorylation by Mec1. Thus, 9-1-1 negatively regulates resection via Rad9 and Rad53. Residual Rad9 binding to chromatin is observed in mutants deficient for 9-1-1 resulting in only partial de-repression of resection at DSBs (57, 63). The positive function of 9-1-1 in resection is through recruitment of the Fun30^SMARCAD1^ chromatin remodeler (64), which counteracts the negative effect of Rad9 on resection (40, 41, 64, 65). In addition, Slx4-Rtt107 competes with Rad9 for binding to γH2A and to Dpb11. In the absence of Slx4-Rtt107, Rad9 recruitment to DSBs is increased and the Rad53 checkpoint is hyper-activated resulting in reduced end resection (48, 66). When more Slx4 is bound to Dpb11 (*rad9*-*2A* mutant) the checkpoint is attenuated resulting in suppression of *sae2Δ* DNA damage sensitivity. We suggest that Dna2-Sgs1 starts resection in *mre11*-*nd* and *sae2Δ* cells generating a recessed 5’ end for 9-1-1 and Dpb11 loading; however, this initial resection is insufficient to displace MRX and Tel1, and when Sae2 is absent, Rad9 accumulates slowing extensive resection by Dna2-Sgs1 and Exo1.

CDK and Mec1/Tel1-catalyzed phosphorylation events play critical roles in regulating resection nucleases. Sae2^CtIP^ is licensed to activate Mre11 endonuclease when CDK activity is high in S and G2-phase cells to ensure end resection occurs when a sister chromatid is available to template homology-dependent repair (12-14); similarly, Dna2 is positively-regulated by CDK-catalyzed phosphorylation (67). Human EXO1 is also activated for end resection by CDK (68). The other positive action of CDK is by phosphorylation of Fun30 and Slx4, which is required for their interaction with Dpb11, and hence to 9-1-1 at the recessed 5’ end (47, 64, 69). While Mec1 and Tel1-mediated phosphorylation of Sae2 is important for resection (70), Mec1 and Tel1 act to repress extensive resection via recruitment of Rad9 and Rad53 to DSBs (71). As described above, Rad9 blocks extensive resection by Dna2-Sgs1 and Rad53 inhibits Exo1 activity. Here, we show that preventing CDK-catalyzed phosphorylation of Sae2 does not impact the checkpoint dampening function of Sae2, consistent with its role in activating Mre11 endonuclease. By contrast, Tel1 and/or Mec1-catalyzed phosphorylation of Sae2 in the vicinity of DSBs provide an additional way to regulate resection by attenuating Rad9 binding and Rad53 activation until resection initiates. This feedback control mechanism activates Dna2-Sgs1 and Exo1 if resection initiation is delayed.

In fission yeast, Mre11 nuclease and Ctp1^Sae2^ are more important for DNA damage resistance than observed in budding yeast (20, 72). As described above, resection in *mre11*-*nd* and *sae2Δ* mutants is highly dependent on Dna2-Sgs1. Langerak et al (22) showed that Rqh1^Sgs1^ barely contributes to resection in fission yeast and instead Exo1 is largely responsible for long-range resection. The poor use of the Dna2-Rqh1 pathway at DSBs in fission yeast could be due to a stronger blockade by Crb2^Rad9^ (73). Although Mre11 nuclease and CtIP are both essential for proliferation of mammary cells, *Ctip*^-/-^ mouse embryos arrest at an earlier stage than Mre11^H129N/H129N^ embryos, suggesting that CtIP, like Sae2 in budding yeast, has functions beyond stimulating Mre11 endonuclease (74, 75).

## MATERIALS AND METHODS

### Media, growth conditions and yeast strains

Rich medium (1% yeast extract; 2% peptone; 2% dextrose, YPD), synthetic complete (SC) medium and genetic methods were as described previously (76). CPT or MMS was added to SC or YPD medium, respectively, at the indicated concentrations. For survival assays, 10-fold serial dilutions of log-phase cultures were spotted on plates with no additive or the indicated amount of drug and incubated for 3 days at 30°C. Diploids heterozygous for relevant mutations were sporulated and tetrads dissected to assess synthetic genetic interactions. Spores were manipulated on YPD plates and incubated for 3-4 days at 30°C. The yeast strains used here, listed in Table S1, are derived from W303 corrected for *RAD5*, and were constructed via crosses or by one-step gene replacement using PCR-derived DNA fragments. To generate the SSA assay system, we modified the BIR assay system described by Donnianni and Symington (77). A 5’ truncated *lys2* fragment was inserted 20 kb strain telomere proximal to a 3’ truncated *lys2* cassette and HO cut site, such that the *lys2* fragments have the same polarity on the left arm of Chr V in a strain expressing *GAL*-*HO*. *RAD51* was deleted from the SSA strains to prevent break-induced replication. Details of plasmid constructions are in the supplementary methods.

### Chromatin immunoprecipitation (ChIP) and co-immunoprecipitation (IP) assays

Yeast cells were cultured in YP containing 2% lactate (YPL) or 2% raffinose (YPR) to log phase and arrested at G2/M phase by adding nocodazole (15 μg/ml) to the medium. For ChIP experiments, cells were collected 90 minutes or 180 minutes after adding galactose to 2% for HO endonuclease induction. After formaldehyde cross-linking and chromatin isolation, Mre11, Rad9-HA, HA-Tel1, Sae2-MYC or γH2A (ab15083, Abcam) were immunoprecipitated as described previously using Mre11 polyclonal antibodies from rabbit serum, anti-HA antibodies (16B12, BioLegend), anti-MYC antibodies (9E10, Santa Cruz) or, respectively (54, 78). Quantitative PCR was carried out using SYBR green real-time PCR mix (Biorad) and primers complementary to DNA sequences located 0.2 or 1 kb from the HO-cut site at the *MAT* locus. Reads were normalized to DNA sequences located 66 kb from HO-cutting site. For Co-IP experiments, cells were collected 1 h after galactose addition for Sae2-MYC induction for no DNA damage control and collected 3 h after MMS (0.1%) treatment or 2 h after Zeocin treatment (30 μg/ml). Rad53 was immunoprecipitated from the cell extracts with anti-Rad53 antibodies (IL-7, gift from M. Foiani), then immunoblotted with IL-7, anti-HA antibodies, anti-MYC antibodies to recognize Rad9-HA and Sae2-MYC, respectively.

### Western blots

Yeast cells were grown to 10^7^ cells/ml in YPD, then CPT, MMS or zeocin was added at the indicated final concentration for 60 min. Cells were released into fresh YPD medium and collected at the indicated time points for TCA precipitation. Cells were resuspended in 0.2 ml 20% TCA and then mechanically disrupted for 5 minutes using glass beads. Beads were washed twice with 0.2 ml 5% TCA each and pellets collected by centrifugation at 3000 rpm for 10 minutes. The pellet was resuspended in 0.15 ml SDS-PAGE sample buffer and proteins separated by SDS-PAGE. Anti-Rad53 antibodies, anti-HA antibodies and anti-MYC antibodies were used for immunoblots.

### SSA assay

Cells containing the SSA reporter were grown for HO induction as described above. Aliquots of cells were removed prior to HO induction (0 h), and at one or two hour intervals after addition of galactose to the media for isolation of genomic DNA. Genomic DNA was digested with EcoRV and the resulting blots hybridized with a PCR fragment corresponding to *LYS2* sequence by Southern blotting. SSA efficiency can be also measured by qPCR. We designed primer pairs to amplify sequences 3 kb downstream of the HO cut site (HOcs) between two *LYS2* homologies (3K_DS), an 3.2 kb upstream of the HOcs (3.2K_US). The Ct values for each primer pair were normalized to ADH1, and the SSA product was calculated by the ratio of 3K_DS/3.2K_US.

## ACKNOWLEDGEMENTS

We thank M.P. Longhese, B. Pfander and M. Smolka for yeast strains and plasmids, and M. Foiani for anti-Rad53 antibodies. This study was supported by grants from the National Institutes of Health (P01CA174653 and R35 GM126997).

---

**Figure S1.**
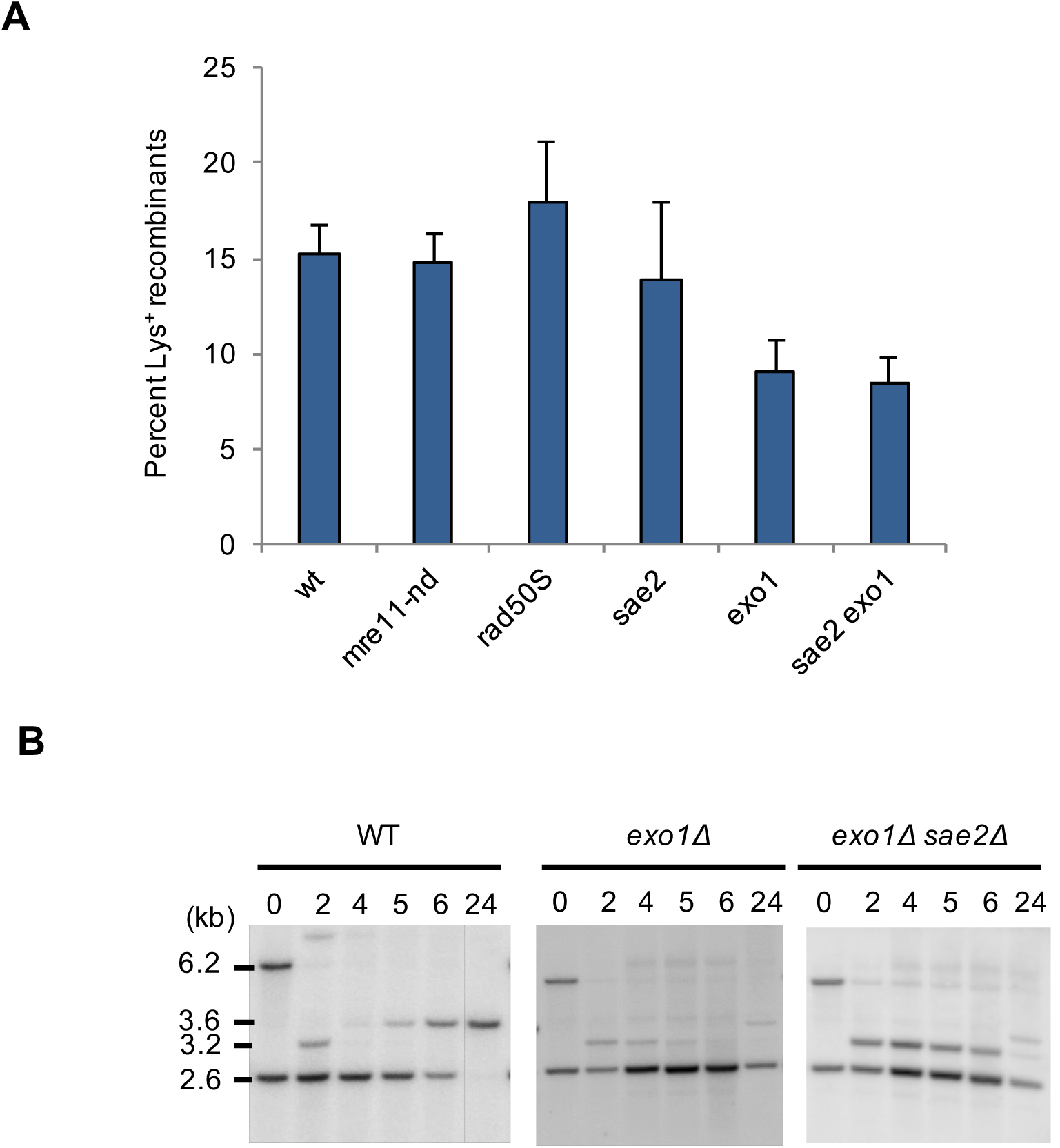
Defects in resection initiation do not influence SSA efficiency. **A** The percent SSA as determined by the ratio of colony forming units (cfu)on SC-LYS + GAL medium to cfu on SC GLU-containing medium for the indicated strains. **B** Southern blot to detect the SSA product in WT, *exo1Δ* and *exo1Δ sae2Δ* strains.

**Figure S2.**
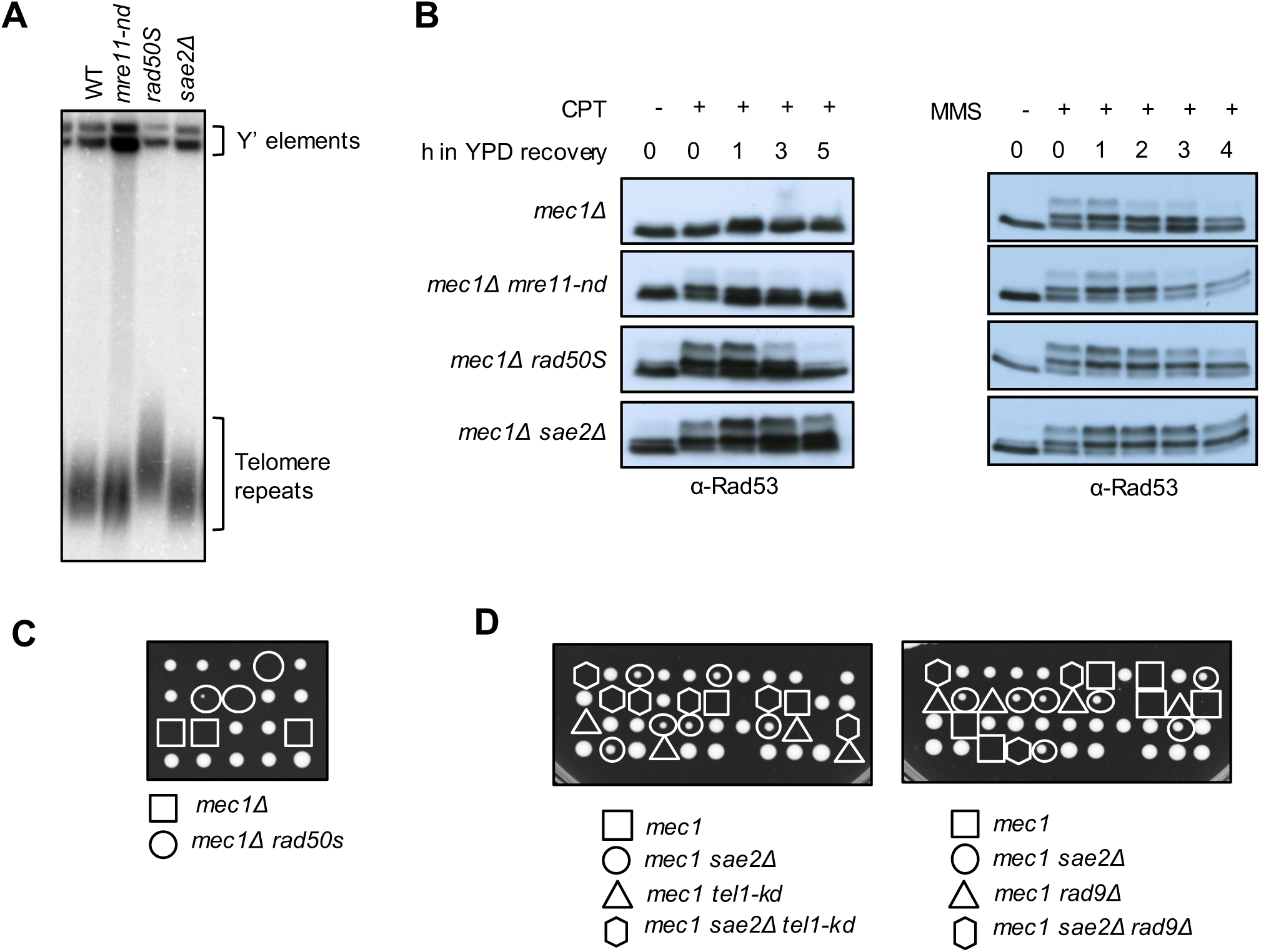
Tel1 hyper-activation in *radSOS* and *sae2Δ* cells. **A** Southern blot of XhoI digested genomic DNA probed with a Y’-telomere probe. **B** Western blot analysis showing Rad53 phosphorylation pattern in response to CPT or MMS. Log phase growing cells (t=0) from indicated strains were treated with 10 μg/mL CPT or 0.015% MMS for 1h and released into fresh YPD (t=0-5). Protein samples from different time points before and after drug treatment were analyzed using anti-Rad53 antibodies. **C** Spore colonies from dissection of a *mec1Δ/MEC1 sml1Δ/SML1 RAD50/rad50S* heterozygous diploid. **D** Spore colonies from dissection of *mec1Δ/MEC1 sml1Δ/SML1 SAE2/sae2Δ TEL1/tel1*-*kd* and *mec1Δ/MEC1 sml1Δ/SML1 SAE2/sae2Δ RAD9/rad9Δ* heterozygous diploids.

**Figure S3.**
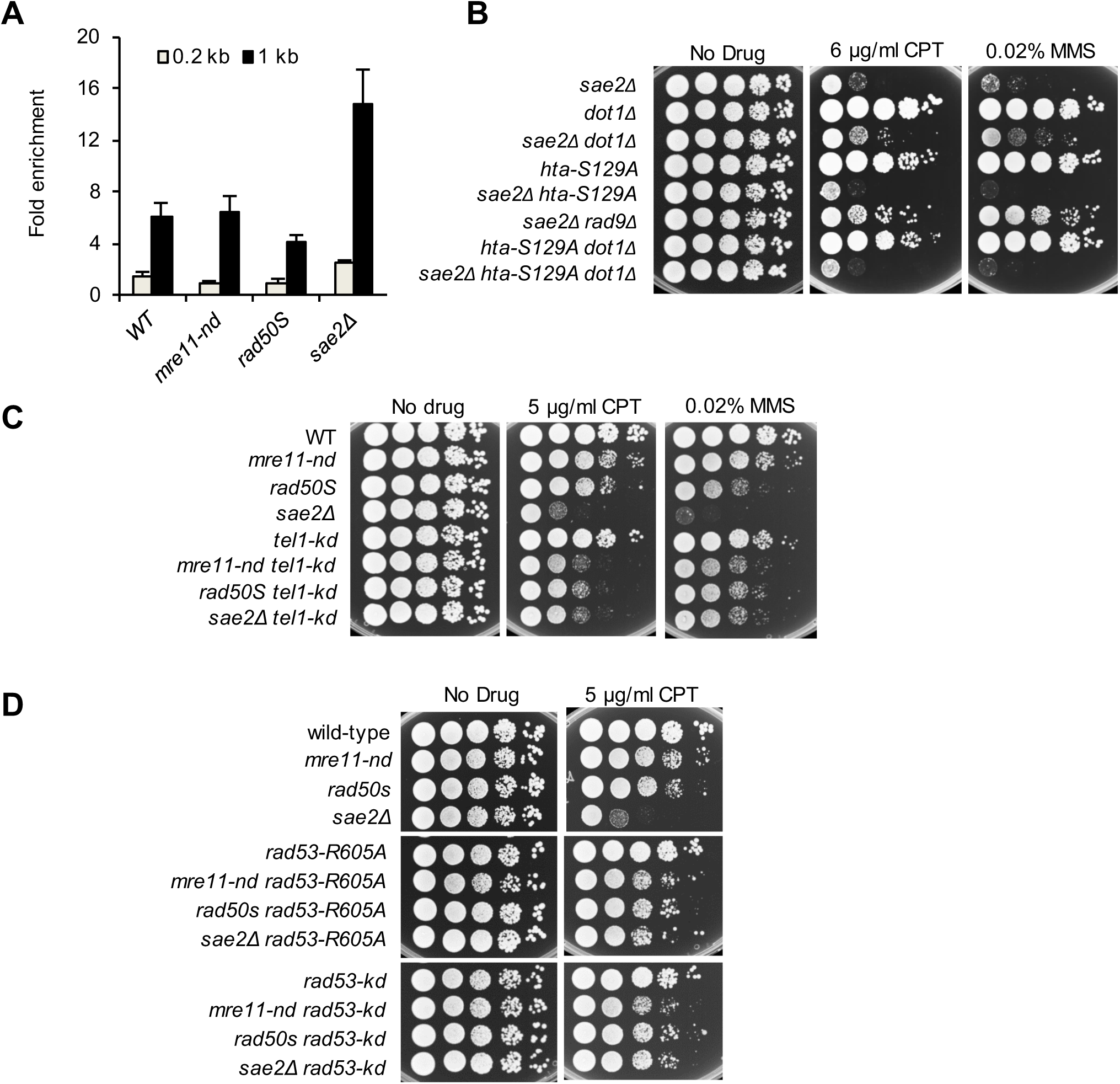
Rad9 chromatin binding and Rad53 kinase activity contribute to *sae2Δ* CPT sensitivity. **A** The relative fold enrichment of γH2A at 0.2 and 1 kb from the HO site was evaluated by ChIP-qPCR after 3 hours HO induction. **B** Ten-fold serial dilutions of *dot1Δ* and *hta*-*S129A* derivatives spotted on plates without drug, or plates containing CPT or MMS at the indicated concentrations. **C** Ten-fold serial dilutions of *tel*-*kd* derivatives spotted on plates without drug, or plates containing CPT or MMS at the indicated concentrations. **D** Ten-fold serial dilutions of *rad53*-*R605A* or *rad53*-*kd* derivatives spotted on plates without drug, or plates containing CPT at the indicated concentrations.

**Figure S4.**
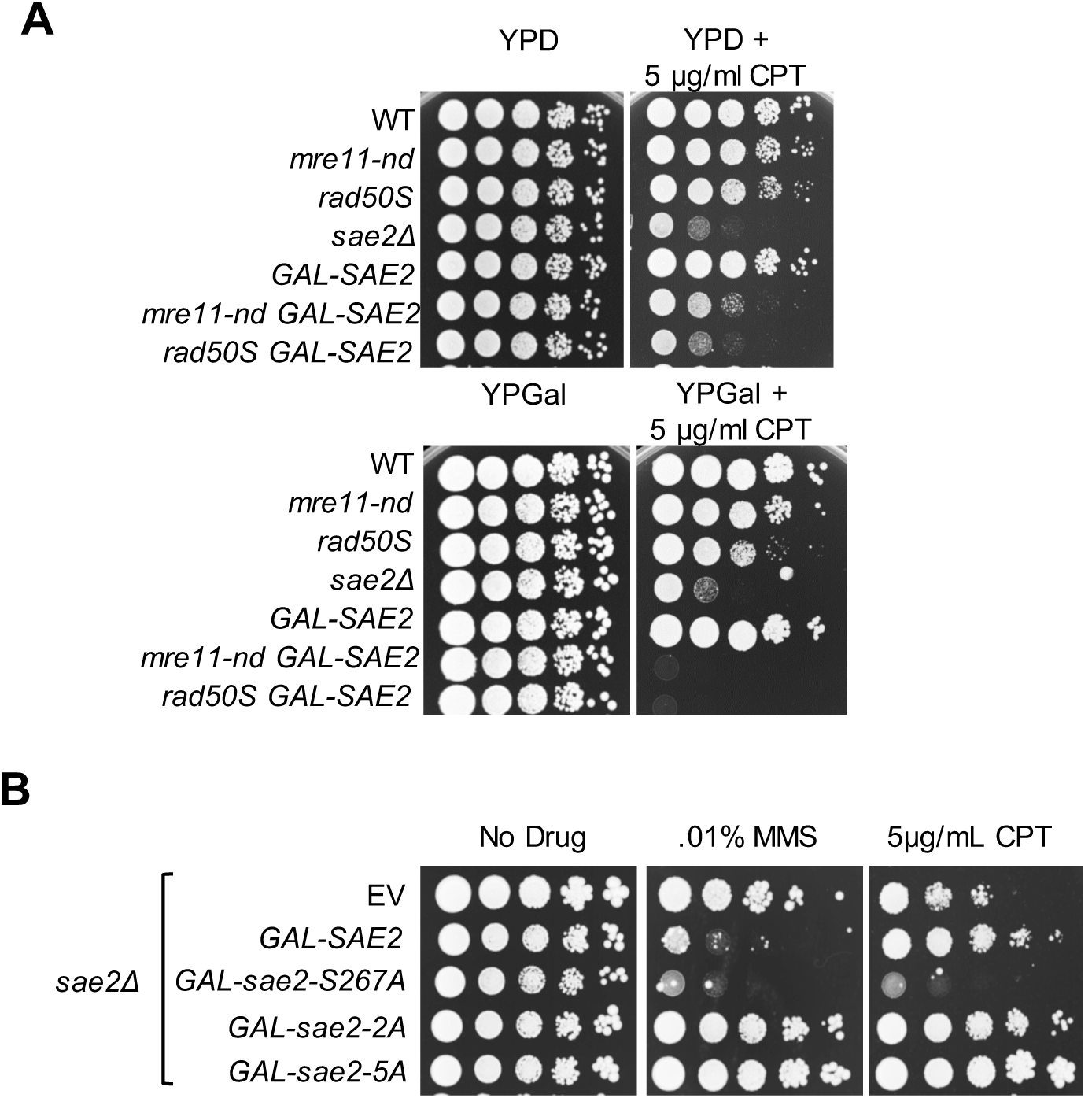
*SAE2* OE is toxic to cells. **A** Ten-fold serial dilutions of indicated strains spotted on YP with glucose -/+CPT (upper panel), or YP with galactose -/+ CPT (lower panel). **B** Ten-fold serial dilutions of *sae2Δ* cells with EV, or the indicated *SAE2* allele expressed from the *GAL* promoter of a low copy number plasmid spotted on YPGal, YPGal + MMS or YPGal + CPT.

## Supplementary Methods

### Plasmids

To construct plasmids pRS426-*SAE2* (2μ) and pRS416-*SAE2* (*CEN*), the 1,038 bp *SAE2* ORF flanked by 370 bp upstream promoter and 36 bp synthetic MYC epitope were amplified by PCR from genomic DNA of LSY0678 and the resulting product was cloned into BamHI-SalI digested pRS426 or pRS416 (primers are available on request). pRS416-*sae2*-5A was made using plasmid pML487 as a template (1). pRS416-*sae2*-S267A and pRS416-*sae2*-T90A, T279A (2A) were made by side-directed mutagenesis of pRS416-*SAE2* (Agilent QuickChange II kit). pRS416-*SAE2*-13MYC was made by PCR amplification of *SAE2* with a 13MYC tag from genomic DNA of LSY3942-8C and the resulting fragment was cloned into NruI-SalI-digested pRS416-*SAE2*. Plasmids pRS416-*sae2*-S267A-13MYC, pRS416-*sae2*-2A-13MYC and pRS416-*sae2*-5A-13MYC were made by sub-cloning NruI-BsiWI fragments containing the phosphorylation site mutant alleles from pRS416-*sae2*-S267A, pRS416-*sae2*-2A, pRS416-*sae2*-5A into NruI-BsiWI-digested pRS416-*SAE2*-13MYC. Sae2 overexpression plasmids, pRS416-*GAL*-*SAE2* was made by PCR amplifying a 693 bp BglII-NruI fragment containing the *GAL1* promoter and the 5’ part of *SAE2* from LSY3942-8C genomic DNA and cloning into BamHI-NruI-digested pRS416-SAE2. Then a SpeI-NruI 699 bp fragments containing the *GAL1* promoter and partial 5’ *SAE2* from *pRS416*-*GAL*-*SAE2* was sub-cloned into SpeI-NruI-digested pRS416-*sae2*-S267A, pRS416-*sae2*-2A, pRS416-*sae2*-5A to obtain pRS416-GAL-*sae2*-S267A, pRS416-GAL-*sae2*-2A, and pRS416-GAL-*sae2*-5A.

**Table S1.**
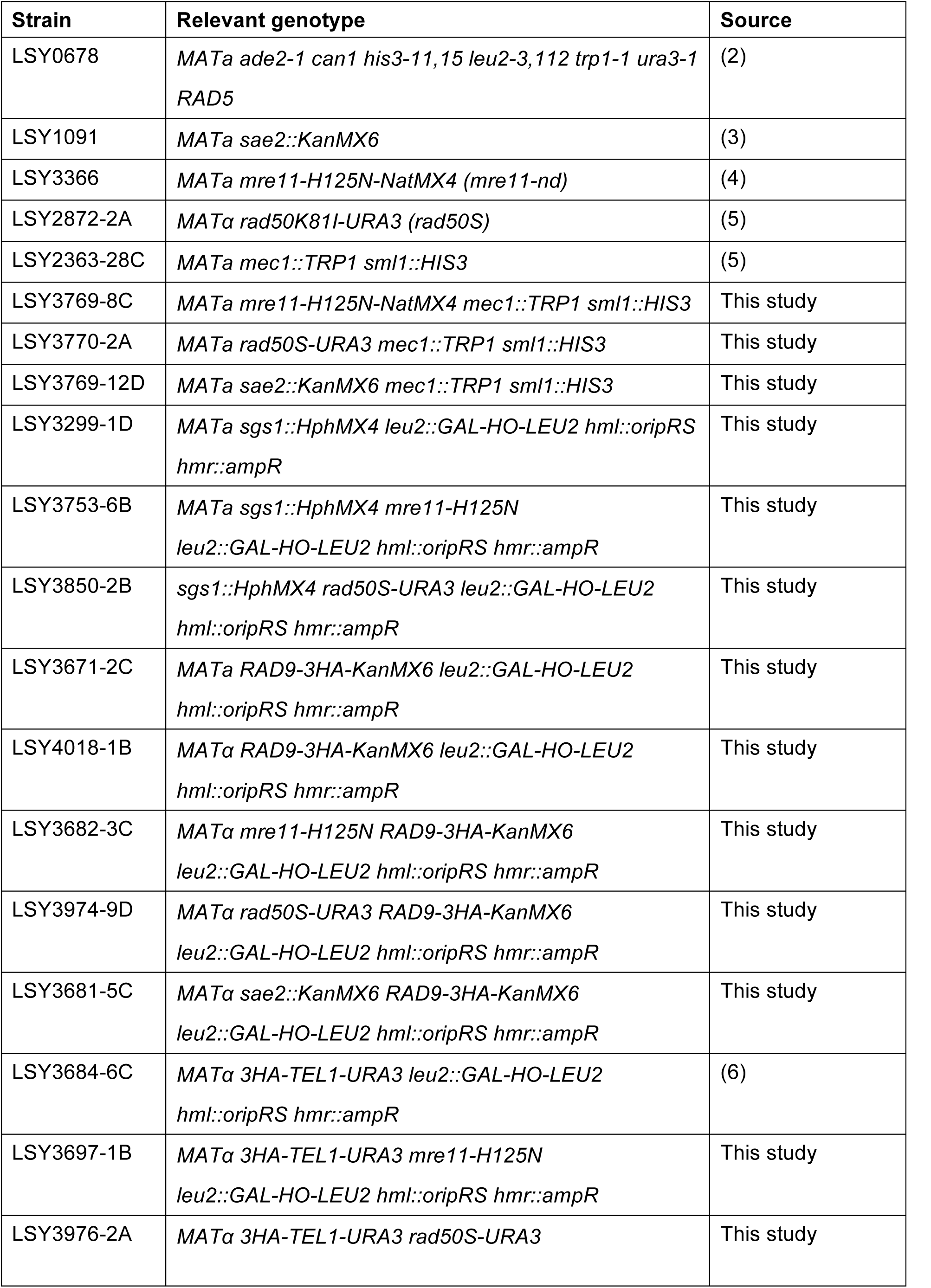
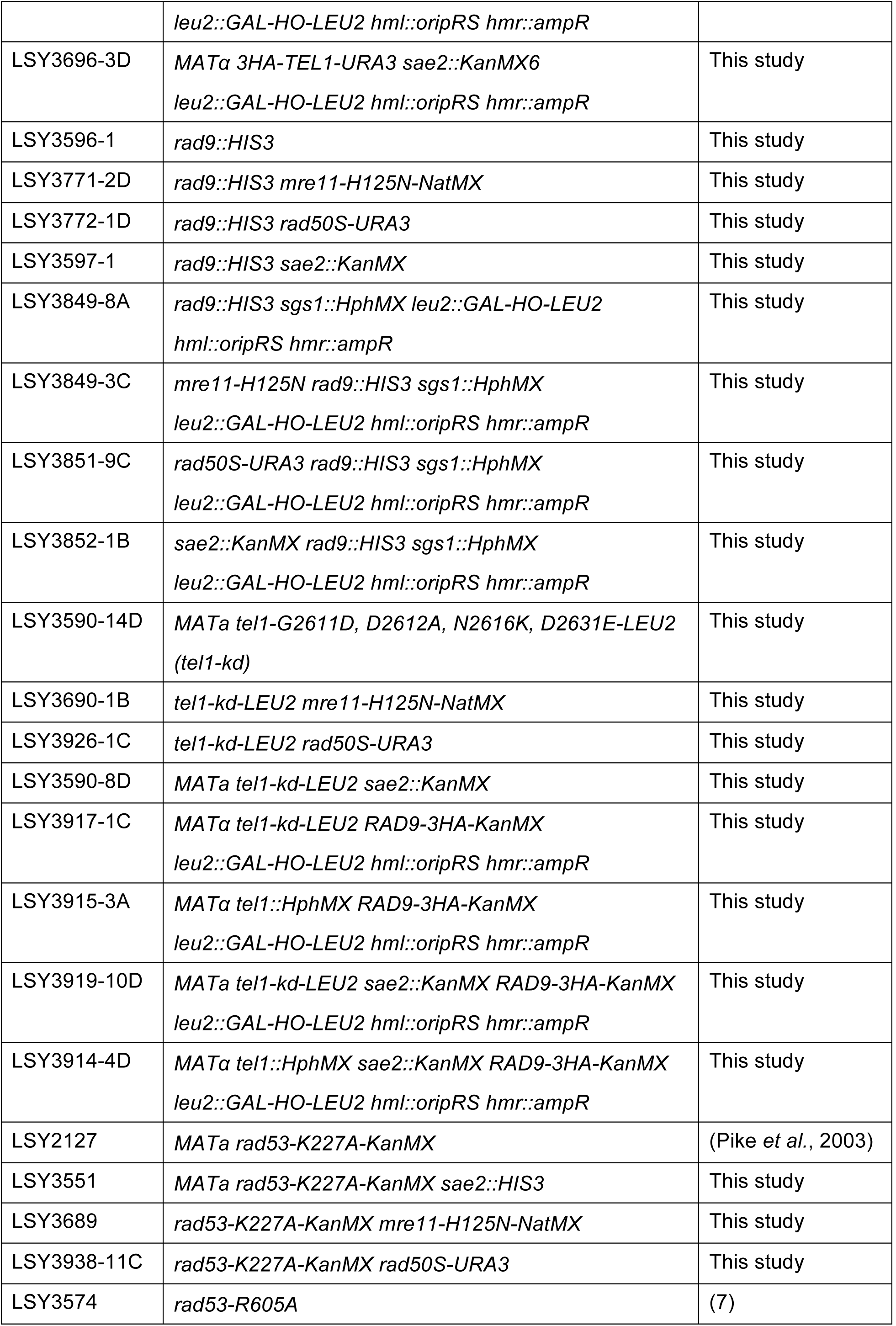
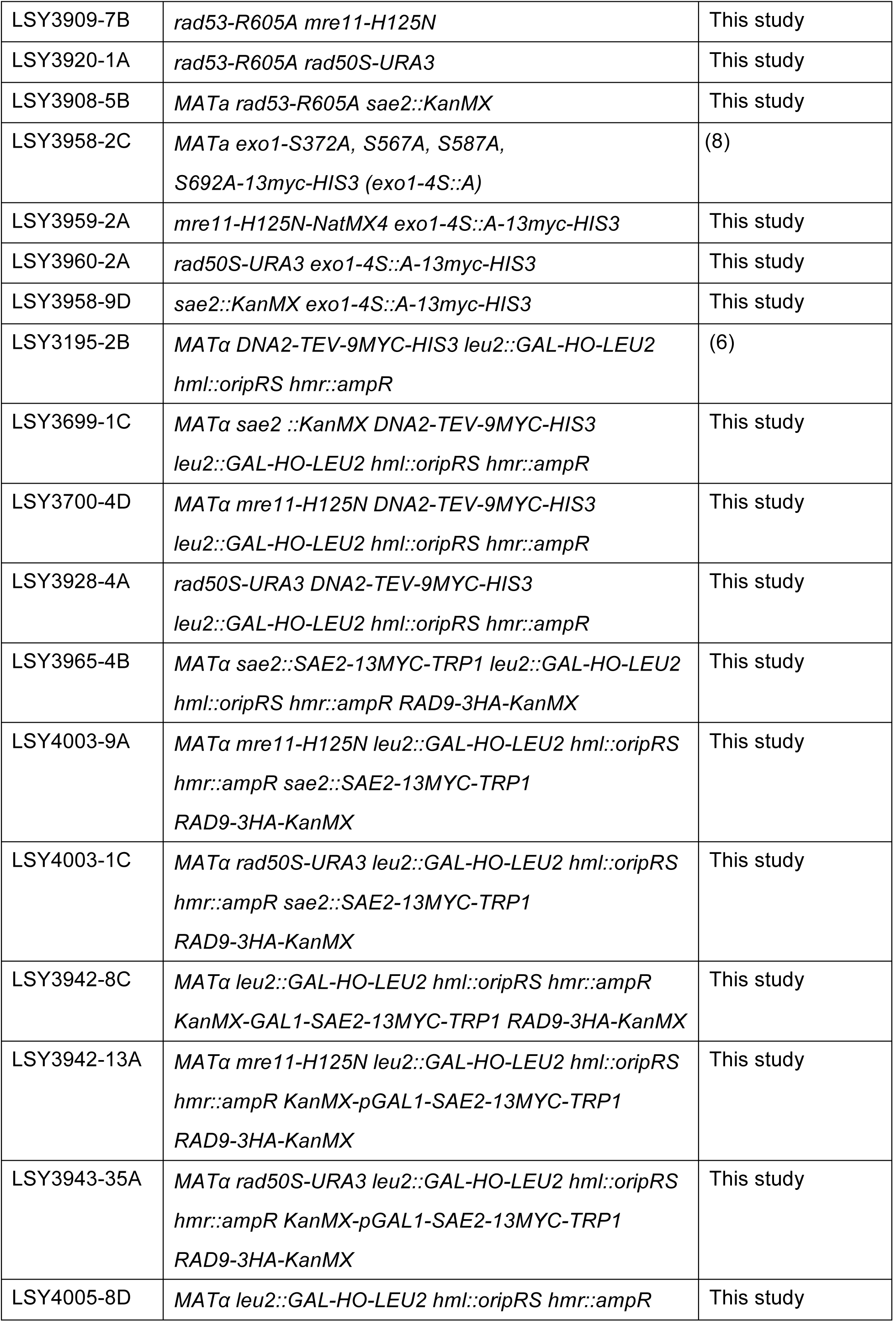
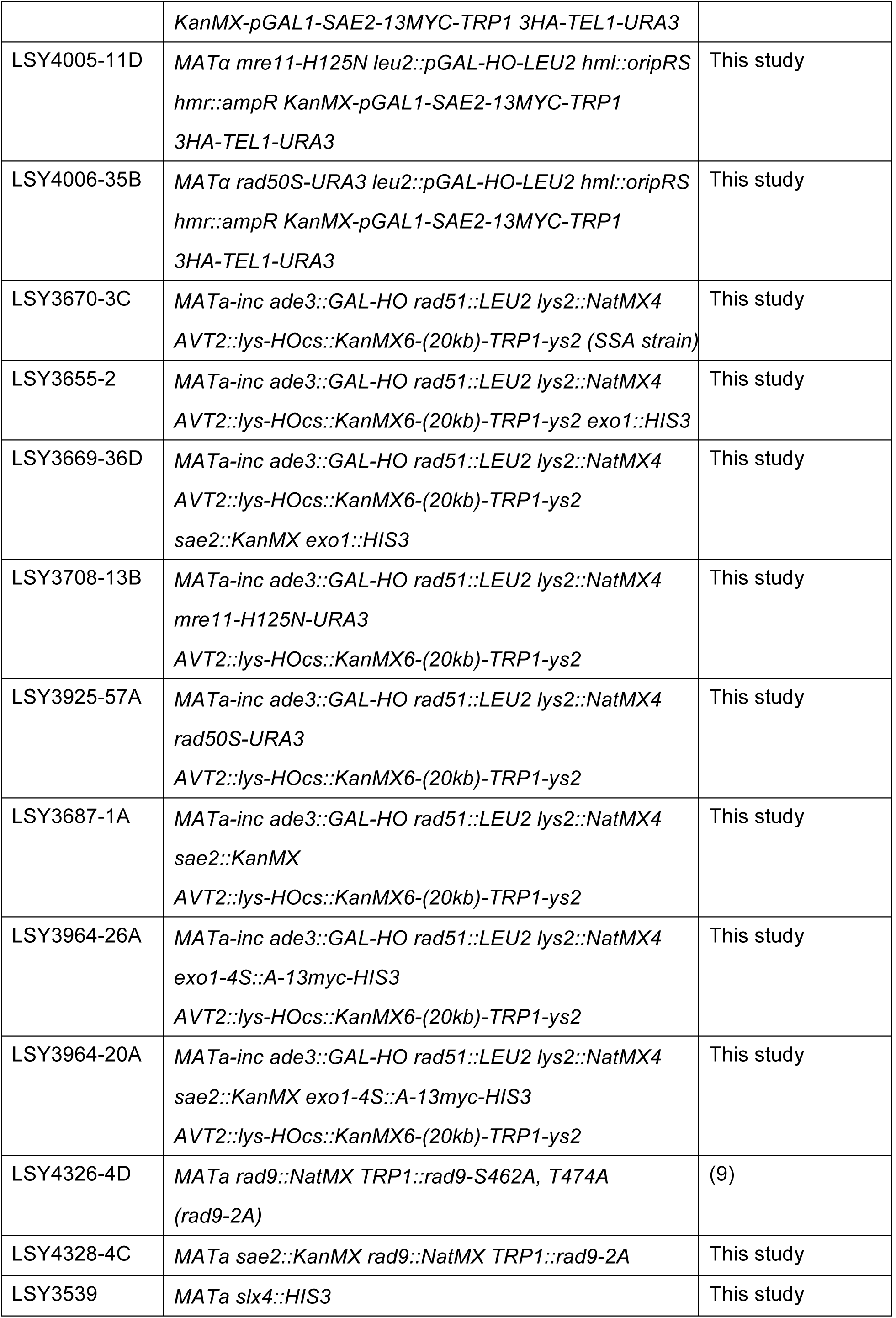
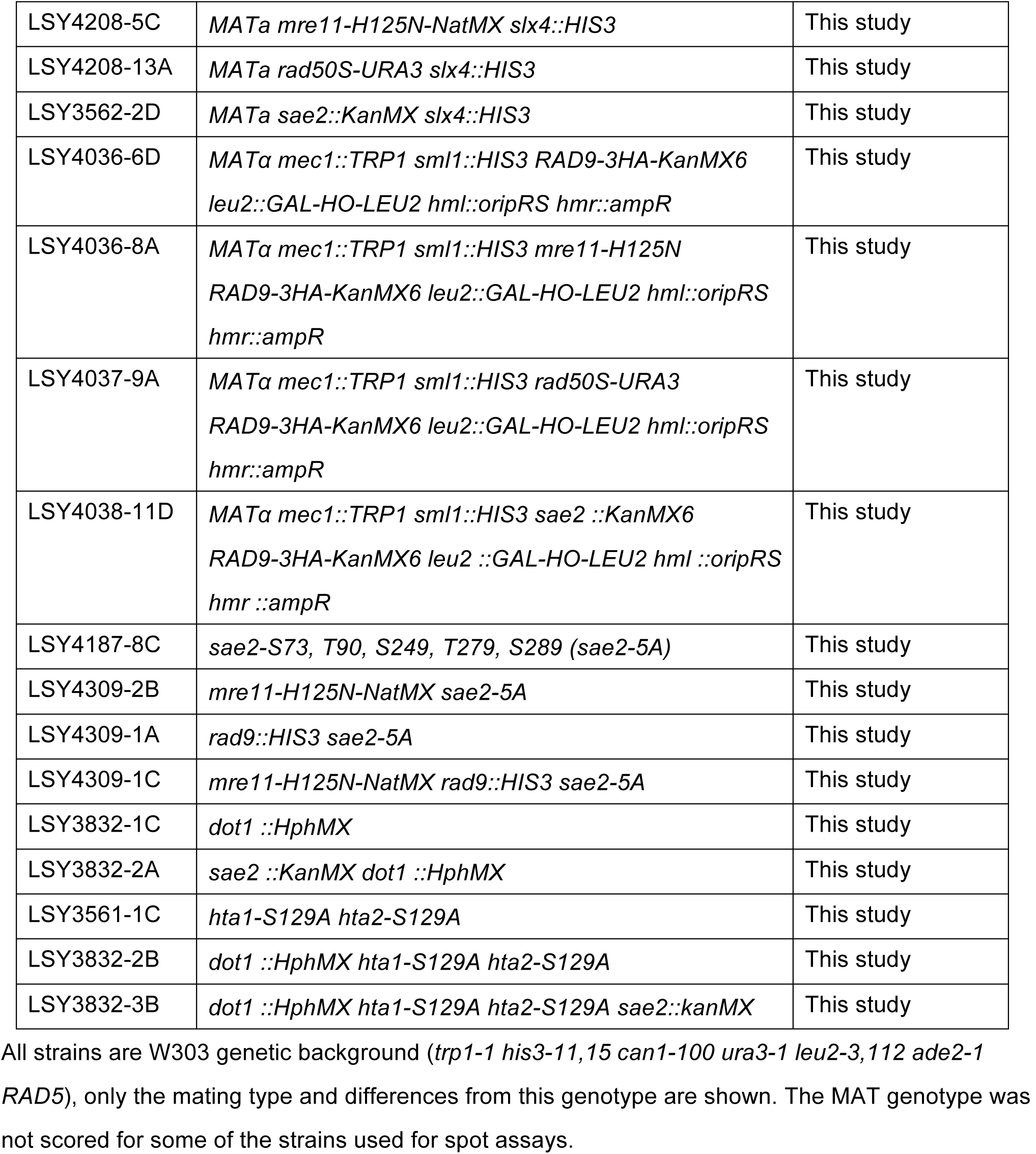
*Saccharomyces cerevisiae* strains used in this study.

